# Longitudinal Prediction of Drug Response in High-Grade Serous Ovarian Cancer Organoid Cultures Aligning with Clinical Responses

**DOI:** 10.1101/2024.05.09.593311

**Authors:** Enrico Cavarzerani, Isabella Caligiuri, Michele Bartoletti, Giuseppe Corona, Tiziana Perin, Antonio Palumbo, Vincenzo Canzonieri, Flavio Rizzolio

## Abstract

High-grade serous ovarian cancer (HGSOC) ranks among the most aggressive gynecological malignancies. Its high mortality stems from frequent recurrence post-primary treatments and development of platinum resistance. Since, traditional drug development via animal models is time-consuming and lacks reproducibility, in precision cancer medicine, primary patient-derived organoids (PDOs) offer a solution by replicating disease pathophysiology and expediting drug screening. We developed an expandable HGSOC organoid platform for rapid drug screening and resistance testing. Our study aimed to validate the stability of seven PDO pairs in predicting drug responses over time. Organoids underwent low- and high-passage drug screenings over nine months, involving 21 conventional and FDA-approved drugs and proteomic analyses. Comparison of in vitro outcomes with clinical data confirmed the platform’s predictive capacity. Notably, a PDO with BRCA1 mutation exhibited resistance to Carboplatin and PARP inhibitors, highlighting organoid models’ clinical relevance for novel targeted therapies such as the Peptidylprolyl Cis/Trans Isomerase, NIMA-Interacting 1 (Pin1), a valuable target for HGSOC patients.

## INTRODUCTION

Deadly diseases need more appropriate biological models to develop additional accurate diagnostic and therapeutic approaches. Mouse models have been the most widely utilize and still are in place since have many advantages including a fast reproduction, limited costs, genetic stability and a solid scientific literature [1], [2], [3]. Thanks to that, scientists discovered many fundamental processes that are at the basis of Mendelian and complex diseases including cancer.

High-grade serous ovarian cancer (HGSOC), a highly lethal disease, gained considerable attention in recent times, owing to breakthroughs in identifying the tissue of origin and the development of novel cell lines and animal models [4], [5], [6], [7]. Additionally, genomic analyses have uncovered actionable mutations and alterations, offering potential avenues for improved treatments, although overall survival rates remain unsatisfactory [8], [9], [10], [11] [4]. Consequently, many scientists continue to explore alternative models to study HGSOC, with the aim of making this disease more manageable [5], [12], [13].

Among in vitro models, the use of organoids represent an emerging approach [14]. Pioneering works demonstrated that defined adult stem cells are able to originate the crypt and villus structures with all differentiated cell types of the intestine and embryonic stem cells are able to form a self-organized structure recapitulating the spatial and temporal aspects of early neuronal development [15], [16]. Following, most of the tissues and organs were modeled utilizing the organoid technology including HGSOC [17], [18]. Single epithelial cells derived from fallopian tubes are able to form organoid cultures in 3D extracellular matrix supplemented with Wnt, Notch, EGF, FGF-10 and TGF-b signaling molecules. The cells were able to proliferate and differentiate in secretory and ciliated cells [19]. Genetic modification of both fallopian tube and ovarian surface epithelium mouse organoids are able to generate HGSOC although with different phenotypic/clinical penetrance and latency and unique molecular profiles that respond differently to chemotherapy [12]. Organoids derived from immunocompetent mice with specific genetic alterations showed genotype-dependent similarities in chemosensitivity, secretome, and immune microenvironment and allowed to develop an effective combination treatment for CCNE1-amplified subgroup of HGSOC [20].

In 2018, Hill Sarah J. et al. established thirty-three short-term cultures of HGSOC human organoids to identify targetable DNA damage repair defects [17]. Functional defects in the homologous recombination (HR) pathway have been shown to correlate with sensitivity to poly (ADP-ribose) polymerase (PARP) inhibitors, while deficiencies in replication fork protection have been associated with sensitivity to carboplatin, CHK1 inhibitors, and ATR inhibitors [17]. The HR pathway plays a crucial role in repairing DNA double-strand breaks, and its dysfunction can lead to genomic instability and tumorigenesis. PARP inhibitors exploit the concept of synthetic lethality in cancer cells with HR deficiencies, leading to cell death [21]. In contrast, replication fork protection mechanisms are essential for maintaining genome stability during DNA replication, and their impairment can sensitize cells to DNA-damaging agents, such as carboplatin, CHK1 inhibitors, and ATR inhibitors [17], [21], [22]. The ability of organoids to predict drug sensitivity was confirmed and expanded demonstrating that it is possible to derived long-term cultures of ovarian cancer (OC) organoids representing all main subtypes. Intra- and interpatient heterogeneity was detected and acquisition of chemoresistance in recurrent disease [23]. These data were confirmed by other groups demonstrating the potential use of organoids for clinical decision [24], [25], [26]. Even our group demonstrated that HGSOC organoids can be used to test drug on the market or to develop novel drugs [27], [28], [29], [30].

While some published papers have supported the utilization of patient-derived organoids (PDOs) as a promising predictive biomarker in cancer treatment, [31], there are a few examples that demonstrate the ability of organoids to predict drug response after long-term culture. Colorectal cancer organoids were tested with 11 different drugs at the beginning and after 1 month of culture with good to fair reproducibility [32]. While some studies demonstrate reproducibility and retention of patient-specific characteristics in OC organoids over long-term cultures [23] [24] [33] [34], none have directly assessed changes in drug response. This highlights a significant gap in current research, emphasizing the need for further exploration into the stability of drug responsiveness in organoid cultures over extended periods. Moreover, we explored the potential to develop novel targeted therapies and testing new inhibitor of Peptidylprolyl Cis/Trans Isomerase, NIMA-Interacting 1 (Pin1), an emerging target of HGSOC as demonstrating by our group [33] [23] [34] [24] [35], [36], [37].

## RESULTS AND DISCUSSION

### Establishment of a Living Biobank of OC Patients

As depicted in **figure 1A, B**, tumor organoid cultures were obtained from seven different patients. All these patient-derived tumor organoids (PDTOs) were effectively cultured from tumor specimens, resulting in organoids that are highly expandable and sustainable for long-term culturing. Two tumor organoids were derived from the primary tissue and 5 from ascites. Of note, organoid formation efficiency did not significantly differ between ascites and primary tissues [38].

**Figure 1.**
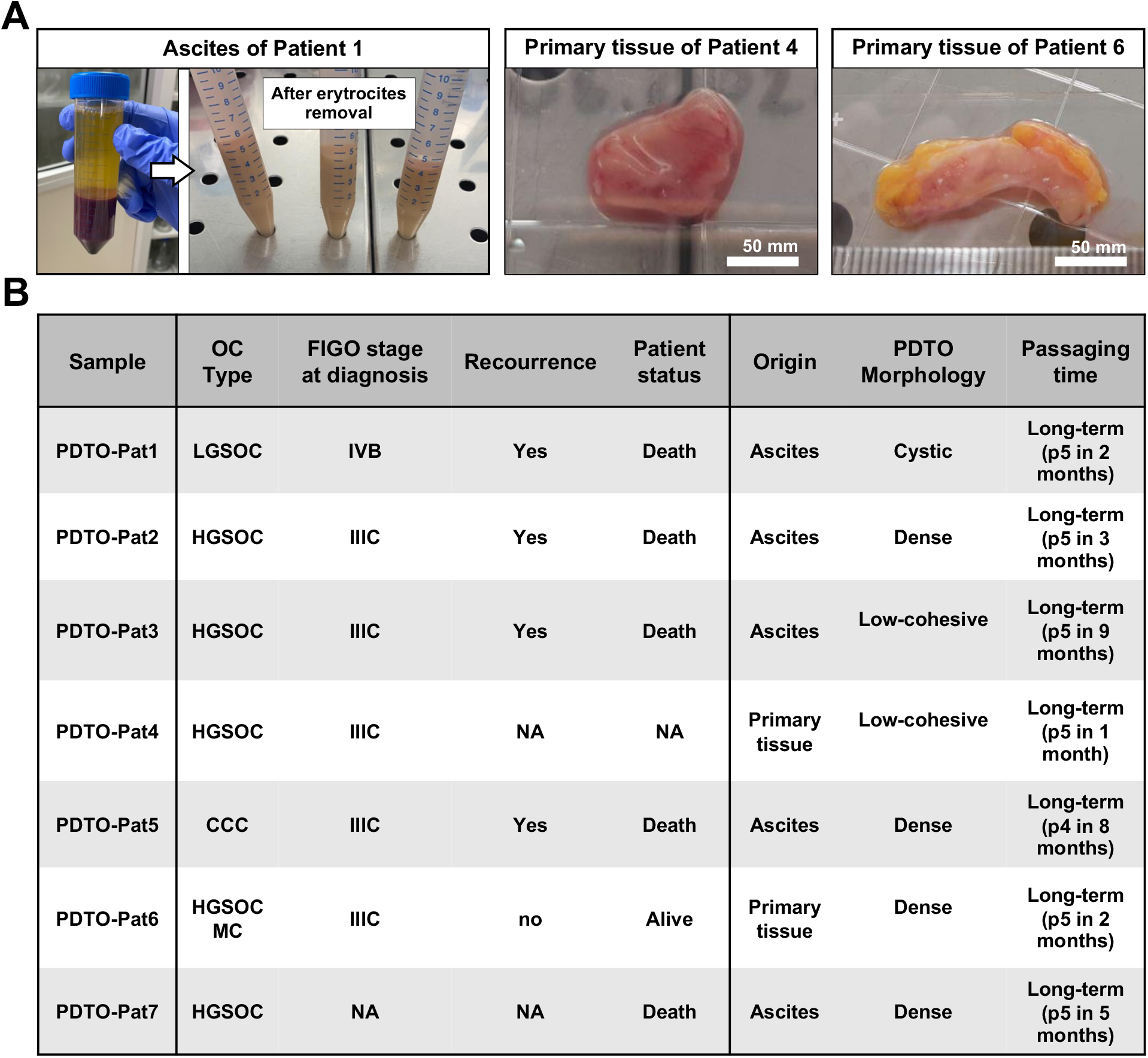
EOC Patients, Establishment of Organoid Lines and biobanking: (**A**) An example of primary tumor and ascites samples derived from patients 1, 4 and 6, before processing for organoid culture. (**B**) Summary of the patient/organoid characteristics: HGSOC, high grade serous ovarian cancer; LGSOC, low grade serous ovarian cancer; CCC, clear cell carcinoma; MC, mucinous carcinoma; long-term cultures refer to organoids with a high passageability (higher than 5 passages).

All patients were diagnosed and treated for FIGO stage IIIB-IV epithelial OC and main histological subtypes and molecular subgroups were well represented.

Patient 1 was diagnosed with low grade serous OC at age of 32. The clinical course showed an aggressive and platinum resistant behavior. Ascites was collected at the time of platinum-resistant recurrence. Molecular analysis performed on formalin fixed, paraffin embedded tumor tissue showed all-RAS-BAF wild type, BRCA 1-2 wild type, mismatch repair proficiency. Patient 2 was diagnosed with FIGO stage IIIC HGSOC and a pathogenic BRCA1 mutation was unveiled. After primary surgery and platinum chemotherapy, she received maintenance with olaparib for only 6-months before experiencing a recurrence of peritoneal disease. Ascites were collected at the time of recurrence. Patients 3, 4 and 7 with HGSOC were treated with cytoreductive surgery and platinum chemotherapy and have a BRCA 1-2 wild type status. Patient 5 was diagnosed with clear cell carcinoma ad she died few months after the end of platinum-based chemotherapy. Patient 6 was diagnosed with HGSOC with a partial mucinous differentiation.

### EOC-Derived Organoids Reproduce the Original Tumor Phenotype

As depicted in **figure 1B** “passaging time”, variations in the growth rate of the tumor organoids under investigation were observed. Notably, achieving passage 5 took one month for some organoids, while it required nine months for others, probably reflecting the heterogeneous composition of the tumors. In this respect, all the organoids have been characterized by hematoxylin and eosin (H&E) staining together with immunohistochemistry (IHC) of PAX8, WT1, CA125 and TP53 markers (**Fig. 2**), evaluated by two pathologists and summarized in **figure 2C**. PAX8 (paired box) gene is a transcription factor primarily associated with fallopian tube development and is not essential for ovarian development. In contrast to PAX8, PAX2 is typically downregulated early in the progression of serous OC. However, PAX8 maintains ubiquitous expression, underscoring its role in processes such as migration, invasion, proliferation, cell survival, stem cell maintenance, and tumor growth as analyzed in the study of Hardy et al. [39]. As a result, PAX8 is utilized as a marker for the detection of HGSOC, with PAX8 expression observed in both primary tumors and ascites, as well as in PDTOs [39]. The expression of Wilms’ Tumor 1 (WT1) has been documented in various adult cancers [40], [41]. Within the female genital tract, its localization is employed as a discriminative marker for distinguishing serous OC from other cancer types. A well-established association between WT1 expression and an unfavorable prognosis in OC is well-documented [42], [43]. Specifically, WT1 has been found to be expressed in 71.4% of Low-Grade Serous OC (LGSOC) and in 57.1% of HGSOC [42]. The serum levels of CA125 are a valuable indicator for anticipating responses to chemotherapy, as well as for assessing the risk of relapse and disease progression in patients with OC [44]. Although it exhibits limited sensitivity in detecting stage I disease and specificity, CA125 is recognized as a predictive marker for preinvasive OC [45], [46]. Elevated CA125 levels are also associated with a higher likelihood of recurrence or disease progression [47]. To compare organoids to their original tumors, we performed H&E staining on neoplastic tissue sections and evaluated the expression of PAX8, WT1, CA125, TP53 and PIN1 (potential) proteins as OC biomarker (**Fig. 2A**). The examined markers were quantified as shown in **figure 2B**, and no statistically significant differences in terms of expression were demonstrated between the primary tumor/ascites and PDTOs. As known, it is therefore possible to assert that the PDTOs closely resemble the structural and molecular characteristics of the tumors from which they originate.

**Figure 2.**
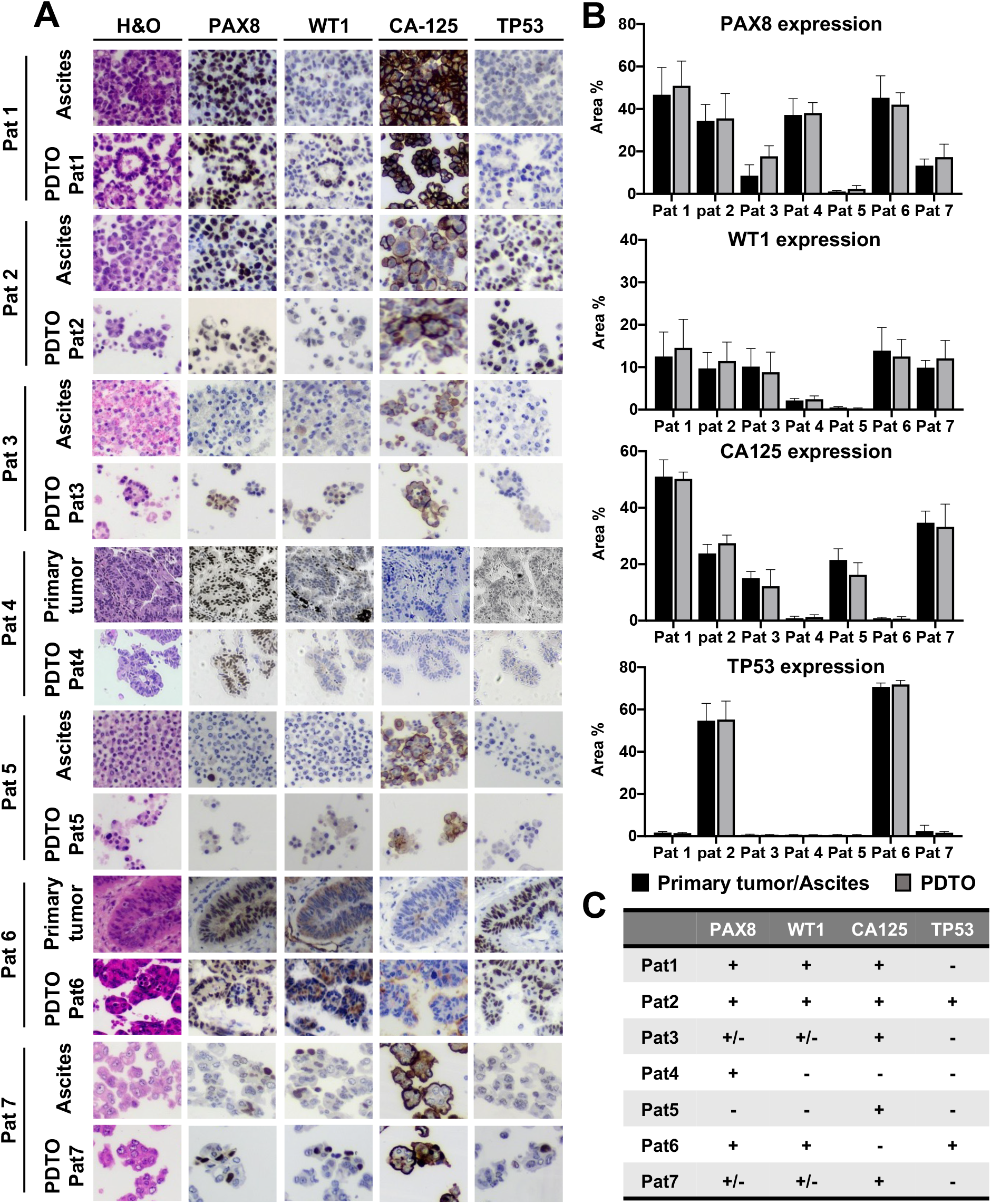
Organoids recapitulate the structure and characteristics at morphological and molecular levels of the tumors from which they are derived: (**A**) H&E staining and IHC of ascites derived from HGSOC (Patient 2, 3, 7), LGSOC (Patient 1), advanced CCC (Patient 5) and HGSOC primary tumor tissues (Patient 4, 6). CA125 (cancer antigen 125), WT1 (Wilms’ Tumor 1), PAX8 (Paired box gene 8) and TP53 were done. (**B**) The percentage area of marker expression, as determined by IHC, was calculated in both primary tissue/ascites and PDTOs. The bar graph illustrates the mean and SD. The p-value was computed using the “RM one-way ANOVA” test; (**C**) summary table of the positivity or negativity of PDTOs to the IHC markers.

### Validating Organoid Stability in Long-term cultures

All organoids in the study were expanded and comprehensively characterized to confirm their fidelity in representing the originating tumor tissue. Subsequently, these PDTOs were cryopreserved to establish a biobank dedicated to EOC organoids. Biobanks play a critical role in preserving and providing access to these valuable organoid specimens for research and medical purposes. They often include various types of organoids, such as those derived from different tissues or disease conditions, and they are crucial for advancing fields like regenerative and personalized medicine. In an organoid biobank, organoids undergo multiple cycles of freezing and thawing. Therefore, it is crucial to establish a model that remains stable over time, especially when considering personalized therapies. Prior to assessing clinical correlations with patients, experiments were conducted to validate the reproducibility of these organoids in both short and long-term cultures. In parallel, morphological features, growth rates, proteomic profiles, and drug screening responses were evaluated in both high and low-passage organoids to ensure that drug responses remain consistent over time.

#### PDTOs morphology, growth and proteomic profile

The 3D organoid structure is formed within 2-4 days of culturing and displays inter-patient morphological heterogeneity. As reported by Maenhoudt et al., the physical structure of organoids varied among samples from patients with epithelial OC, showing distinct characteristics [26], [48], [49]. As shown in **figure 3A** some exhibited a compact and “dense” structure with little to no central cavity (organoids derived from Patient 2, 5, 7), while others had a chaotic arrangement characterized by weak cellular bonds, termed “low-cohesive” (organoids derived from Patient 3, 4, 6). Additionally, a subset presented a “cyst-like” structure (Patient 1), defined by a thin layer of cells surrounding a substantial central space, similar to the findings reported by Kopper et al. [23]. Also, the viability was measured over time (9 days) and there were no statistically significant differences between the growth rate of low- and high-passage organoids (**Fig. 3B**). The maintenance of the structural morphology as well as the growth rate over time suggest that the phenotype remains conserved over time.

**Figure 3.**
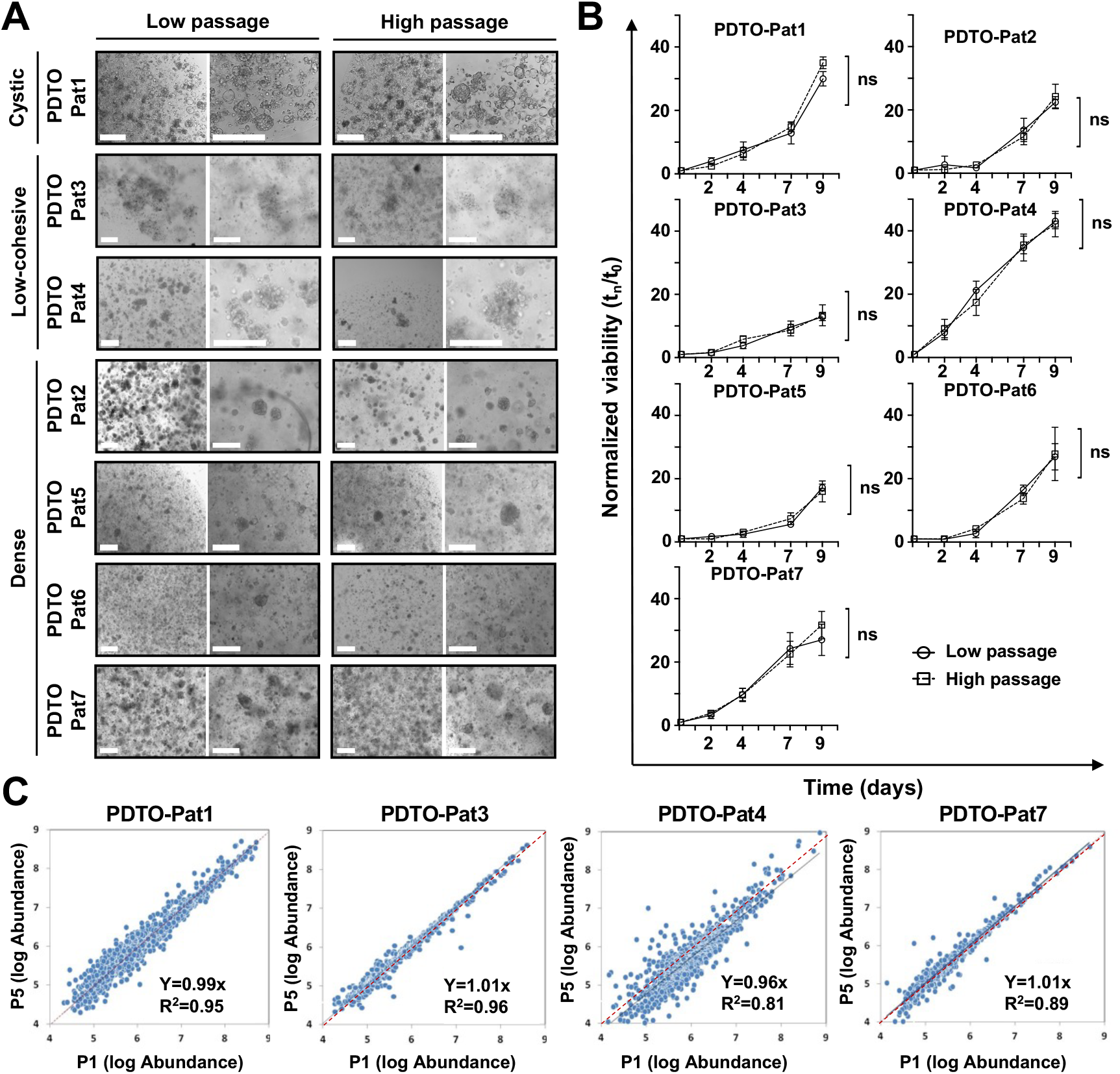
Conservation of morphological and proteomic profiles in PDTOs over time: (**A**) Different morphology of OC organoids. Representative bright-field images of organoid cultures at low and high passages (Short and Long-term expansion). Scale bar is 200 μm. (**B**) Growth curve of the PDTOs. Cell viability was measured using PrestoBlue HS Assay, mean ± SD is reported, and p-values were calculated using a two-tailed Student’s t-test. * p ≤ 0.05, ** p ≤ 0.01, *** p ≤ 0.001 and **** p ≤ 0.0001. (**C**) Scatterplot of the protein abundance of 4 PDTOs (PDTO-Pat1, 3, 4, and 7) at low and high passages. The abundance of every protein is calculated as a triplicate and shown as a dot in the graph. The coefficient of regression was between 0.81 and 0.96, while the slope was between 0.96 and 1.01.

A proteomic profile of the organoids derived from patients 1, 3, 4, 7 was established. The abundance of the proteins at low and high passage was measured and compared.

Protein expression at low passage was found to be strongly correlated with expression at high passage (**Fig. 3C**); measurements at low passage were closely correlated with those measured at high passage. More than the 80% of the analyzed proteins do not undergo significant changes indicating that the proteomic profile of organoids remains relatively stable over time. This result is in accordance with the experiment conducted by Kooper et al. [23] in which was evaluated the genomic landscape copy number variations (CNV) between early and late passage HGSOC organoids [23]. Their findings revealed that CNVs were remarkably preserved even after extended passages, thereby confirming that organoids maintain a consistent protein expression profile even at later stages.

#### A drug screening profile in short- and long-term organoid cultures

Finally, the response of PDTOs to drugs was assessed in both short-term and long-term cultures. PDTOs were treated with serial dilution concentrations of selected drugs for 96 hours, and cell viability was determined to obtain a dose-response curve. From this dose-response curve, the area under the curve (AUC) and IC50 values (for effective drugs) were calculated. As shown in **figure 4A**, which represents an example of drug treatments on PDTOs of patient 7, the dose-response curve exhibited no statistically significant changes in the treatments in both short-term and long-term organoid cultures, regardless of drug efficacy. This naturally extends to the IC50 and AUC values, as exemplified by PDTO-Pat7 in **figure 4B and C**, where no statistically significant differences were observed between drug treatments in low and high-passage PDTOs. This experiment was systematically conducted for all drugs and all PDTOs examined in the study. As depicted in **figure 4D and 4E**, a linear correlation of IC50 and AUC values was observed, calculated after treatment of short-term and long-term PDTO cultures. These data confirm what was reported by van de Wetering et al, who demonstrated that, for certain drugs, the AUC remained unchanged after treatment of colon PDTOs at different cellular passages [32].

**Figure 4.**
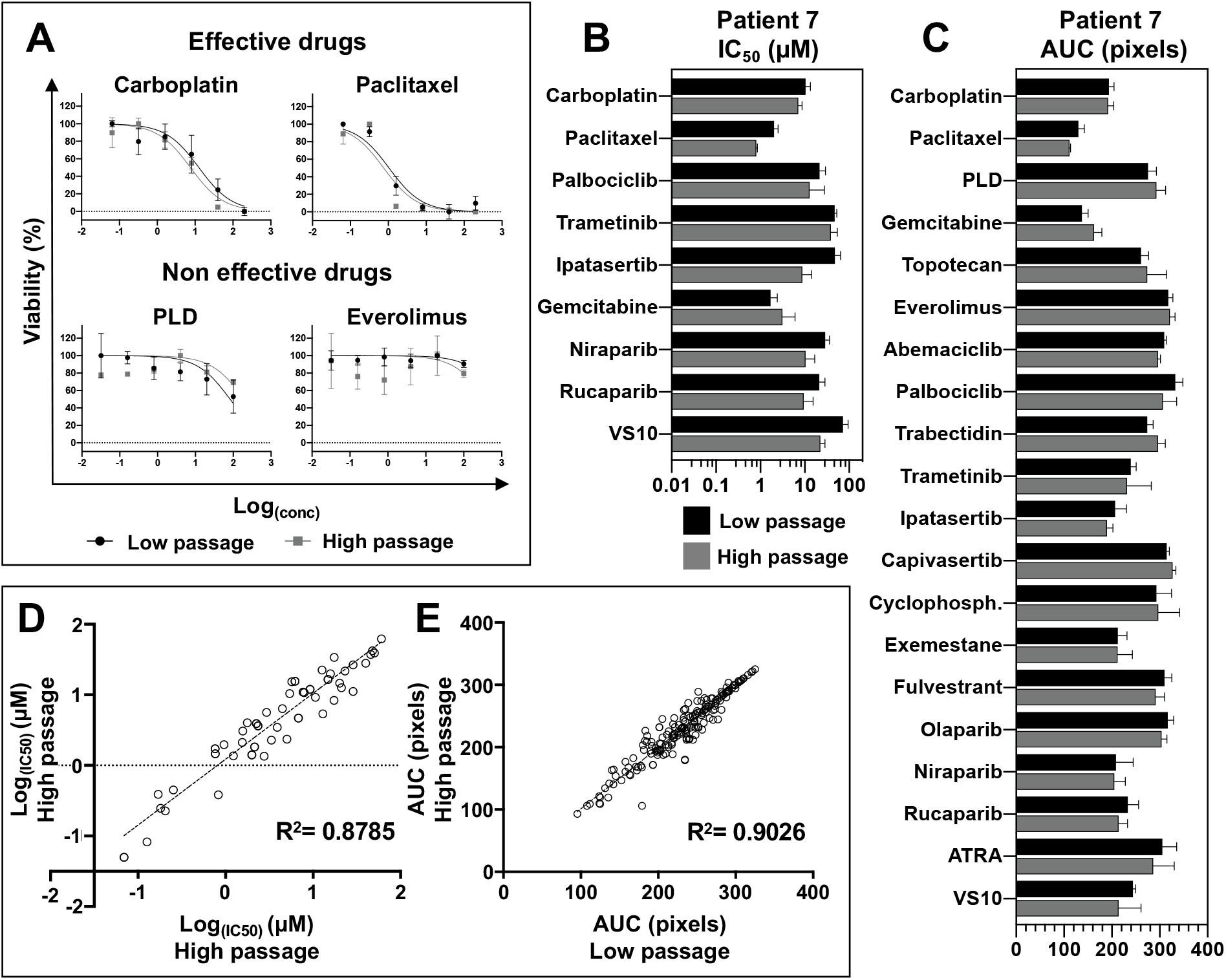
PDTOs maintain the ability of predict patient response to drugs over time: (**A**) Comparison between high and low passage dose-response curves of the same drugs: dots represent the mean and error bars represent the SD of quintuplicates. (**B**) Reproducibility of drug response profiles for effective drugs in PDTO-Pat7 at low and high passages. Bars represent IC50 values from 5 replicates, with SD reported. (**C**) Reproducibility of drug response profiles for effective drugs in PDTO-Pat7 at low and high passages. Bars represent AUC values from 5 replicates, with SD reported. Scatterplot illustrating the correlation of log_(IC50)_ values (**D**) and AUC (**E**) at low and high passages for all the PDTOs. Each point represents a replicate of 5 values. The linear correlation coefficient, calculated as the Pearson coefficient, is reported in the figure.

### Drug response correlation among ex-vivo PDTOs and patients

The PDTOs were treated with the selected drugs for a duration of 96 hours, allowing us to generate dose-response curves from which we derived the IC50 values (**Fig. 5a**). The IC50 values were used to create the heatmap as shown in **figure supplementary 1** and **figure 5b**. This approach enabled us to differentiate between PDTOs that were resistant and those that were sensitives to the drugs used in the study, even discerning different degrees of sensitivity, as described in **Fig. 5b** (blue/green “effective drug”, red “very effective drug”). Subsequently, the in vitro outcome was compared with clinical responses of the patients, establishing the clinical relationship as outlined below.

**Figure 5.**
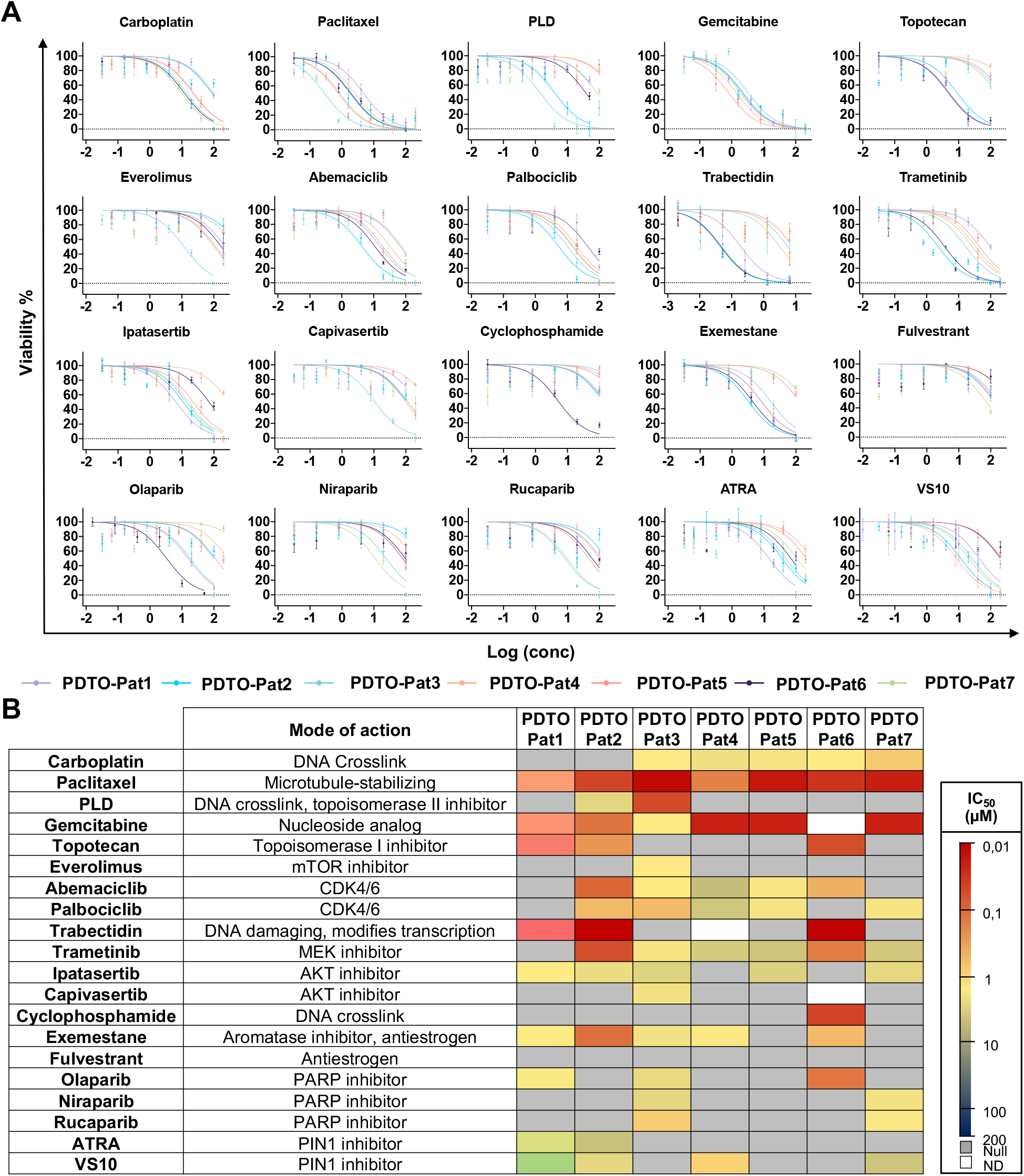
OC organoids as a platform for drug screening: (**A**) Dose–response curves of the PDTOs treated with the compounds. Dots represent the mean of the ten replicates (average of the five replicates for low and high passage each’s). Error bars represent the SEM. ** < p = 0.01, N.S.: not significant (one-way ANOVA). Data analyses were performed using the GraphPad Prism 7.0b software. (**B**) A heatmap derived from the IC50 values was designed for all the patients, in particular grey represent “resistance to the drug”, blue/green “effective drug”, red “very effective drug” and white (ND) “non determined”.

The PDTO-Pat1 was derived from ascites after palliative paracentesis for a relapsed platinum resistant LGSOC previously treated with platinum salts, taxanes, antracyclin, tamoxifen and letrozole. According to the real clinical benefit, the PDTO showed resistance to carboplatin, Pegylated Liposomal Doxorubicin (PLD) while it was sensitive to paclitaxel (7-months clinical benefit) and gemcitabine (6-months clinical benefit). These in vitro sensitivities corresponded to in vivo responses of 6 months to weekly paclitaxel and gemcitabine (the longest PFSs experienced by the patients). Interestingly, the PDTO was resistant to trametinib, a targeted agent actually in label for this rare histotype. The patient received trametinib, but it was discontinued for adverse events (nausea and vomiting) and rapid disease progression. Trabectedin and topotecan for which the PDTO showed sensitivity, were not administered in clinic. The second patient was diagnosed with HGS advanced, germline BRCA1 mutated OC. The PDTO-Pat2 was derived from ascites when she experienced a partially platinum-sensitive relapse after primary platinum-based chemotherapy and maintenance with olaparib for 6 months. According to the clinical benefit, the PDTO was resistant to carboplatin and olaparib, as for other available PARP inhibitors. This result is noteworthy given the potential utility of PDTO even in the presence of strong and clinically validated predictive biomarker as the BRCA mutation. In fact, the clinical benefit from olaparib was unusually short (6 months) even in the presence of a BRCA mutation and this was in line to the resistance to Olaparib observed in the PDTO. The third patient showed a high spectrum of responsiveness to compounds, in particular platinum salts, taxanes and PARPi. This patient was not amenable for upfront or interval cytoreduction and was addressed for a primary platinum-based chemotherapy with a normalization of CA125 (from 100 UI/ml to 23 UI/ml). Maintenance with niraparib was started after a complete biochemical response for 6 months when the patient experiences a disease recurrence. She died few months later. The fourth patient performed the primary cytoreduction in our Institute and then was referred in a near-home hospital for medical therapy. The PDTO was obtained from tumor tissue and no data regarding clinical response to therapies was obtained. The fifth patient was diagnosed with advanced clear cell carcinoma. After primary therapy, the patient experienced a platinum-sensitive recurrence and at that time, the PDTO was obtained from ascites. The in vitro sensitivity to platinum corresponded to the platinum-free interval of 12 months seen in clinic while resistance to other drugs observed in vitro corresponded to the known chemoresistance of this histotype. The sixth patient was diagnosed with a mixed histology high grade serous and mucinous intestinal-like OC. She was treated with first line platinum-paclitaxel and then maintenance with bevacizumab. The clinical benefit and responsiveness to platinum correspond to the platinum sensitivity of the PDTO derived from primary tumor ascites. The patient is alive and disease-free. The seventh patient was diagnosed with HGSOC at the age of 90 years. She experiences a clinical benefit from single agent carboplatin, but she had a disease progression after the 5^th^ administration.

It is noteworthy that the in vitro drug response data derived from PDTOs align precisely with the clinical data obtained from patients, demonstrating a 100% correlation. This robust concordance underscores the profound relevance of this model within the domain of personalized therapy. It effectively anticipates drug resistance, thereby proffering alternative therapeutic strategies with the potential to significantly extend patient survival.

### Pin1 as an innovative target in OC therapy

In addition, during the drug screening, it was possible to test Pin1 inhibitors and compare them with drugs commonly used in the treatment of OC. Our research group has demonstrated, as reported by Russo Spena et al. that Pin1 is overexpressed in HGSOC patients and knocking down Pin1 reduces tumor cell growth in vitro, and in vivo [37]. This suggests that Pin1 could serve as an innovative target for OC.

IHC analysis show that Pin1 expression is similar in primary tumors and derived PDTO **(Fig. 6A, B)** and is overexpressed in 4 out of 7 PDTO **(Fig. 6B)**. Interestingly, Pin1 inhibitors have shown effectiveness only in PDTOs that exhibit Pin1 overexpression **(Fig. 6C)**. Specifically, ATRA was effective in PDTO1 and PDTO2, while VS10 showed efficacy in PDTO1, PDTO2, PDTO4, and PDTO7 (**Fig. 6C**). These results are not surprising since, as previously reported by Russo Spena et al. [35], VS10 is a more potent Pin1 inhibitor compared to ATRA. These findings are in line with the results obtained by Cavarzerani et al. who developed a passive microfluidics platform to do drug screening in PDTOs of HGSOC in which 2 out of 5 PDTOs of the study were sensitive to ATRA [36]. Furthermore, in line with findings from Saorin et al. [50], Pin1 inhibitors appear to exhibit an expression-dependent effect, as they are effective only in PDTOs where PIN1 is overexpressed (PDTO1, 2, 4, and 7).

**Figure 6.**
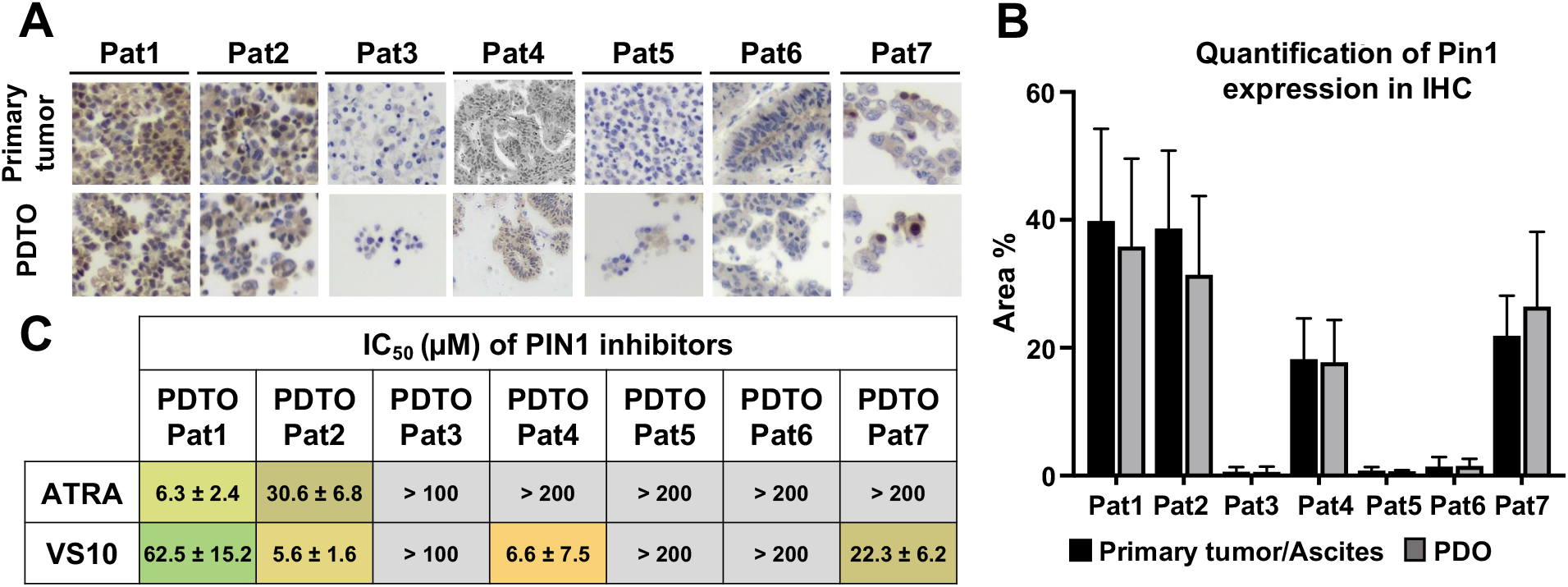
Effectiveness of Pin1 Inhibitors in PDTOs with PIN1 Overexpression: (**A**) Immunohistochemistry of PIN1 in ascites derived from HGSOC (Patient 2, 3, 7), LGSOC (Patient 1), advanced clear cell carcinoma (Patient 5) and HGSOC primary tumor tissues (Patient 4, 6) (**B**) The percentage area of marker expression, as determined by IHC, was calculated in both primary tissue/ascites and PDTOs. The bar graph illustrates mean of 7 replicates, with SD. The p-value was computed using the “RM one-way ANOVA” test; (**C**) IC50 values (μM) of ATRA and VS10 are reported. Data were obtained from four replicates. The numbers represent the mean and SD.

Notably, in some cases (PDTO1 and PDTO2), Pin1 inhibitors were effective in patients resistant to Carboplatin, which is the leading therapy for ovarian cancer (**Fig. 5)**.

## MATERIAL AND METHODS

### Organoid cultures

Organoids were derived from completely de-identified specimens; however, written informed consent for research purposes was obtained to collect the samples at the Pathology Unit of the National Cancer Institute (CRO) of Aviano. For ascitic fluids, cells in the pellets were isolated by centrifugation at 1,000 rpm for 10 minutes and subsequently washed twice with Hank’s Balanced Salt Solution (HBSS, Gibco, Massachusetts, United States). To remove erythrocytes from the fluid, cold red blood cell lysis buffer (Roche Diagnostics, Basel, Switzerland) was added and incubated on ice with continuous stirring for 10 minutes. The cell pellet was centrifuged at 1,000 rpm for 10 minutes and resuspended in Cultrex RGF BME, Type 2 (Bio-techne, Minnesota, USA). Solid tumor tissues were incubated in Dulbecco’s Modified Eagle’s Medium/Nutrient Mixture F-12 Ham without Phenol red supplemented with Levofloxacin (100 μg/mL), Vancotex (25 μg/mL), Ciproxin (5 μg/mL), Gentamicin (200 μg/mL), and Fungizone (5 μg/mL) for 30 minutes. The tissues were then finely minced into pieces of approximately 0.5–1 mm diameter using fine dissection scissors. A solution of 4 mg/mL collagenase IV (Gibco, Massachusetts, United States) was added to the minced tissues, which were incubated at 37°C for no more than 45 minutes and mechanically dissociated by pipetting. The cell clusters were pelleted by centrifugation at 1,000 rpm for 10 minutes, resuspended in an appropriate volume of Cultrex RGF BME, Type 2 (Bio-techne, Minnesota, United States), and then seeded in 24-well cell culture plates. Following the polymerization of Cultrex, 500 μL of organoid culture medium, as described by Kopper et al., was added to each well and replenished every 3 days [23]. The organoids were cultured at 37°C in a 5% CO2 environment.

### Immunohistochemistry

Sections of formalin-fixed, paraffin-embedded ascites and solid tumor PDO were used for histopathological analyses. Organoids were recovered from BME using ice-cold cell recovery solution (Corning, New York, United States) in accordance with manufacturing protocols, fixed in phosphate-buffered 10% formalin, and embedded in 500μL of Bio-Agar (Bio-Optica Milano Spa, Milano, ITA). Five μm sections were stained with H&E using a Leica ST5020 multistainer and 2μm sections were cut for IHC analysis. The IHC was performed with an UltraVision LP Detection System HRP DAB kit (Thermo Scientific, Waltham, USA). Heat-induced antigen retrieval was performed using 10 mM citrate buffer pH 6.0. The following antibodies were used to characterize PDTOs and parental tumors: PAX8 (ProteinTech Group, Germany, EU), WT1 (Abcam, U.K.), CA-125 (Santacruz Biotechnology, TX), TP53 (ProteinTech Group, Germany, EU) and PIN1 (Santa Cruz Biotechnology, California, USA).

### Viability Assays

Clusters of organoids were homogenized in an appropriate volume of Cultrex RGF BME, Type 2, and 2 μL of this homogenate and seeded in 96-well plates. Viability assessments were conducted using the PrestoBlue HS Assay at 2, 4, 7, and 9-day intervals, following manufacturer’s protocols. Viability measurements were performed in four replicates, and p-values were calculated using a two-tailed Student’s t-test with GraphPad Prism (La Jolla, CA, United States). The p-values were denoted as follows: * p ≤ 0.05, ** p ≤ 0.01, *** p ≤ 0.001, and **** p ≤ 0.0001.

### Half-maximal inhibitory concentration (IC50) and deciphering drug effectiveness

Clusters of organoids were mixed in an appropriate volume of Cultrex RGF BME, Type 2, and 2 μL of this mixture were seeded in 96-well plates in five replicates for low and high passage organoids both. After 24 hours the organoids were treated with six different concentrations of the following drugs: Carboplatin, Paclitaxel, PLD, Gemcitabine, Topotecan, Everolimus, Abemaciclib, Palbociclib, Trabectidin, Trametinib, Ipatasertib, Capivasertib, Cyclophosphamide, Exemestane, Fulvestrant, Olaparib, Niraparib, Rucaparib, ATRA and VS10. After 96 h, cell viability was measured using CellTiter-Glo 3D (Promega, Madison, WI, United States). The luminescence was acquired with BioTek Synergy H1. Logistical dose-response curves were used to calculate IC50 and AUC using GraphPad Prism (La Jolla, CA, US).

The interpretation of the IC50 values for the construction of the heatmap was done on the basis of the algorithm proposed by Brook E. et al. [51]. The initial qualitative assessment of the drug response curve is essential to ascertain whether the calculation of the IC50 value is viable, thereby discriminating drugs for which the patient is sensitive from those that are not (shown in the heatmap in **figure 5** in grey).

The IC50 values for drug-sensitive PDTOs were calculated and organized according to their increasing magnitude; a color gradient from red (indicating the lowest IC50 value) to blue (assigned to 200 μM) was assigned. The method used for this interpretation process is exemplified in (**Fig. S1**).

### Proteomic analysis

The PDTOs (approximately 400,000 cells) were isolated and collected using Cell Recovery Solution (Corning, New York, USA) to eliminate the Basement Membrane Extract (BME). They were subsequently subjected to three washes with Hank’s Balanced Salt Solution (HBSS). Peptide extraction was performed using the EasyPep™ MS Sample Prep Kits (Thermo Fisher Scientific Inc, Massachusetts, USA), following the manufacturer’s protocol. Subsequently, the samples were lyophilized, and then reconstituted in 0.1% Formic Acid in MilliQ water. Two mg of desalted digested peptides were loaded and fractionated on Peptide XBridge C18 150 x 2.1 mm column (Waters, Milford, MA, USA) maintained at 40°c and flowed at 0.2 mL/min using Horizon Vanquish UPLC instrument (Thermos Scientific, San Jose, CA, USA). Mobile phase A consisted of water and 0.1% formic acid while mobile phase B consisted of acetonitrile 0.1% formic acid. Peptides are eluted from the column using the following gradient: mobile phase B increased from 1% to 35% over 30 min, followed by 90% B in 5 min, and the column was washed for 1 min at 90% B and then re-equilibrated for 10 min at 1% B. The Eluting peptides from the column were analyzed on a quadrupole-ion trap-Orbitrap QExactive Plus mass spectrometer (Thermo Scientific, San Jose, CA). Orbitrap survey MS1 scans from 375 m/z to 1500 m/z were performed at a resolving power of 70,000 at 200 m/z with an AGC target of 3×106 ions and maximum injection time set to 100 ms. All tandem MS scans were performed on precursors with 2-8 charge states, using HCD fragmentation with normalized collision energy of 20 and dynamic exclusion of 30 s. The instrument was operated in the TNop10 mode and the MS/MS scan was performed at the resolving power of 35,000 using an ion count target set to 1×105 and the maximum injection time of 114 ms. Precursors were isolated using a quadrupole isolation window of 2.2 m/z. Proteome Discoverer 2.5 software (Thermo Scientific, San Jose, CA, USA) processed the raw acquisition file for protein ID and label-free quantification of organoid proteomics profile at early and late stages. The parameters used for ID and LFQ were mass tolerance of 10 ppm for MS1 and 0.02 Da for MS2. The target false discovery rate (FDR) was set to 1% while MS/MS spectra were searched by Sequest against a reviewed Homo sapiens database (SwissProt, November 2023, 20418 entries) using the following parameters: trypsin digestion, maximum of two missed cleavages, cysteine carbamidomethylation as fixed modification, methionine oxidization, protein N-terminal acetylation, and de-methylation as variable modifications. Label-free quantification was performed by exploiting unique and razor peptides for protein abundance calculation.

## CONCLUSIONS

A comprehensive analysis demonstrates that OC organoids maintain original tumor histological characteristics and biomarker expression, such as PAX8, WT1, CA125 and TP53. The organoids and corresponding tumors remained highly similar at the proteomic level, even after extended passaging. This congruence is also evident in their consistent drug response during a drug screening, which remains stable even after prolonged passaging. This is of paramount significance in the context of biobank creation, as the organoids will undergo multiple freeze-thaw cycles, necessitating the establishment of a model that exhibits enduring stability over time.

Furthermore, this PDTO model has demonstrated its capability to replicate the patient’s response to therapeutic treatment in a clinical validation study. An illustration of the significance of this model for personalized therapy is exemplified by the case of PDTO-Pat2, derived from a patient who had a germline BRCA-1 mutation. PARP inhibitors are standard of care in the maintenance setting of high-grade disease in response to platinum therapy. Despite this, about 50% of patients have a disease recurrence and are both platinum and PARP inhibitor resistant, even in the presence of a BRCA 1-2 mutation or a status of homologous recombination deficiency. In this instance, the utilization of a model capable of predicting the patient’s drug response would have provided a useful information for the clinician. This acquires even greater importance in diseases like OC, where in the platinum and PARP inhibitor resistance setting there are currently no validated biomarkers to guide the clinician’s choice among standard single agent chemotherapy. Moreover, PDTO could serve as a platform for new drugs screening to foster the advent of new treatments in clinic.

Finally, this PDTO model has demonstrated the effectiveness of PIN1 inhibitors in a subset of OC patients where PIN1 was overexpressed. It is worth noting that in two cases, these inhibitors were even effective in patients resistant to platinum-based therapy. These data suggest that PIN1 could be a valid and innovative target for OC therapy.

### Disclaimer

This manuscript has undergone grammar and syntax correction using ChatGPT, an AI language model developed by OpenAI. While every effort has been made to improve the clarity and accuracy of the language in this document, the authors acknowledge that the final content and scientific interpretations remain the sole responsibility of the authors and their collaborators. ChatGPT has been used exclusively to enhance the manuscript’s readability and expression, and not to generate any type of data.

## Data availability

The data supporting the findings of this study are available within the paper, supplementary Information files and from the corresponding authors upon reasonable request. Source of data are provided with this paper.

## Author Contributions

E.C. involved in the Conceptualization, Experimental data and Writing

I.C was participated in the data analysis, Review & Editing;

G.C involved in proteomic analysis and Review & Editing

M.B. involved in clinical data analysis and Review & Editing

T.P., A.P contributed in the Biological Samples collection and analyses;

V.C contributed in the Biological Samples analysis, Review & Editing;

F.R. involved in the Conceptualization, data analysis and Review & Editing:

All authors read and approved the final manuscript.

## Funding acquisition

This research was funded by the Ministry of Health, Ricerca Corrente (V.C.); Fondazione AIRC per la Ricerca sul Cancro (Grant AIRC IG23566) (F.R.).

## REFERENCES

[1] W. C. Chiu, D. L. Ou, and C. T. Tan, “Mouse Models for Immune Checkpoint Blockade Therapeutic Research in Oral Cancer,” International Journal of Molecular Sciences 2022, Vol. 23, Page 9195, vol. 23, no. 16, p. 9195, Aug. 2022, doi: 10.3390/IJMS23169195.

[2] D. S. Chulpanova, K. V. Kitaeva, C. S. Rutland, A. A. Rizvanov, and V. V. Solovyeva, “Mouse Tumor Models for Advanced Cancer Immunotherapy,” International Journal of Molecular Sciences 2020, Vol. 21, Page 4118, vol. 21, no. 11, p. 4118, Jun. 2020, doi: 10.3390/IJMS21114118.

[3] E. Li, L. Lin, C. W. Chen, and D. L. Ou, “Mouse Models for Immunotherapy in Hepatocellular Carcinoma,” Cancers 2019, Vol. 11, Page 1800, vol. 11, no. 11, p. 1800, Nov. 2019, doi: 10.3390/CANCERS11111800.

[4] C. Yee, K. A. Dickson, M. N. Muntasir, Y. Ma, and D. J. Marsh, “Three-Dimensional Modelling of Ovarian Cancer: From Cell Lines to Organoids for Discovery and Personalized Medicine,” Frontiers in Bioengineering and Biotechnology, vol. 10. Frontiers Media S.A., Feb. 10, 2022. doi: 10.3389/fbioe.2022.836984.

[5] E. K. Colvin and V. M. Howell, “Why the dual origins of high grade serous ovarian cancer matter,” Nature Communications 2020 11:1, vol. 11, no. 1, pp. 1–4, Mar. 2020, doi: 10.1038/s41467-020-15089-z.

[6] R. Perets et al., “Transformation of the Fallopian Tube Secretory Epithelium Leads to High-Grade Serous Ovarian Cancer in Brca;Tp53;Pten Models,” Cancer Cell, vol. 24, no. 6, pp. 751–765, Dec. 2013, doi: 10.1016/J.CCR.2013.10.013.

[7] J. Kim, D. M. Coffey, C. J. Creighton, Z. Yu, S. M. Hawkins, and M. M. Matzuk, “High-grade serous ovarian cancer arises from fallopian tube in a mouse model,” Proc Natl Acad Sci U S A, vol. 109, no. 10, pp. 3921–3926, Mar. 2012, doi: 10.1073/PNAS.1117135109/SUPPL_FILE/PNAS.201117135SI.PDF.

[8] J. Nair et al., “Resistance to the CHK1 inhibitor prexasertib involves functionally distinct CHK1 activities in BRCA wild-type ovarian cancer,” Oncogene 2020 39:33, vol. 39, no. 33, pp. 5520–5535, Jul. 2020, doi: 10.1038/s41388-020-1383-4.

[9] M. Lupia et al., “Integrated molecular profiling of patient-derived ovarian cancer models identifies clinically relevant signatures and tumor vulnerabilities,” Int J Cancer, vol. 151, no. 2, pp. 240–254, Jul. 2022, doi: 10.1002/IJC.33983.

[10] M. Garziera et al., “Identification of Novel Somatic TP53 Mutations in Patients with High-Grade Serous Ovarian Cancer (HGSOC) Using Next-Generation Sequencing (NGS),” International Journal of Molecular Sciences 2018, Vol. 19, Page 1510, vol. 19, no. 5, p. 1510, May 2018, doi: 10.3390/IJMS19051510.

[11] M. Javellana et al., “Neoadjuvant Chemotherapy Induces Genomic and Transcriptomic Changes in Ovarian Cancer,” Cancer Res, vol. 82, no. 1, pp. 169–176, Jan. 2021, doi: 10.1158/0008-5472.CAN-21-1467/674072/AM/NEOADJUVANT-CHEMOTHERAPY-INDUCES-GENOMIC-AND.

[12] S. Zhang, T. Zhang, I. Dolgalev, H. Ran, D. A. Levine, and B. G. Neel, “Both Fallopian Tube and Ovarian Surface Epithelium Can Act as Cell-of-Origin for High Grade Serous Ovarian Carcinoma,” bioRxiv, p. 481200, Nov. 2018, doi: 10.1101/481200.

[13] D. Hao et al., “Integrated analysis reveals tubal- and ovarian-originated serous ovarian cancer and predicts differential therapeutic responses,” Clinical Cancer Research, vol. 23, no. 23, pp. 7400–7411, Dec. 2017, doi: 10.1158/1078-0432.CCR-17-0638/14700/AM/INTEGRATED-ANALYSIS-REVEALS-TUBAL-AND-OVARIAN.

[14] “JCB: Perspective How do we define organoids and 3D cultures?,” 2017, doi: 10.1083/jcb.201610056.

[15] T. Sato et al., “Single Lgr5 stem cells build crypt-villus structures in vitro without a mesenchymal niche,” Nature, vol. 459, no. 7244, pp. 262–265, May 2009, doi: 10.1038/nature07935.

[16] M. Eiraku et al., “Cell Stem Cell Article Self-Organized Formation of Polarized Cortical Tissues from ESCs and Its Active Manipulation by Extrinsic Signals,” Stem Cell, vol. 3, pp. 519–532, doi: 10.1016/j.stem.2008.09.002.

[17] S. J. Hill et al., “Prediction of DNA Repair Inhibitor Response in Short Term Patient-Derived Ovarian Cancer Organoids,” Cancer Discov, vol. 8, no. 11, p. 1404, Nov. 2018, doi: 10.1158/2159-8290.CD-18-0474.

[18] S. Bose, H. Clevers, and X. Shen, “Promises and Challenges of Organoid-Guided Precision Medicine,” Med (N Y), vol. 2, no. 9, p. 1011, Sep. 2021, doi: 10.1016/J.MEDJ.2021.08.005.

[19] M. Kessler et al., “The Notch and Wnt pathways regulate stemness and differentiation in human fallopian tube organoids,” Nat Commun, vol. 6, Dec. 2015, doi: 10.1038/NCOMMS9989.

[20] S. Iyer et al., “Genetically DEFI ned syngeneic mouse models of ovarian cancer as tools for the discovery of combination immunotherapy,” Cancer Discov, vol. 11, no. 2, pp. 384–407, Feb. 2021, doi: 10.1158/2159-8290.CD-20-0818/333548/AM/GENETICALLY-DEFINED-SYNGENEIC-MOUSE-MODELS-OF.

[21] D. Lemaçon et al., “MRE11 and EXO1 nucleases degrade reversed forks and elicit MUS81-dependent fork rescue in BRCA2-deficient cells,” Nat Commun, vol. 8, no. 1, Dec. 2017, doi: 10.1038/s41467-017-01180-5.

[22] Z. Baranski et al., “Aven-mediated checkpoint kinase control regulates proliferation and resistance to chemotherapy in conventional osteosarcoma,” Journal of Pathology, vol. 236, no. 3, pp. 348–359, Jul. 2015, doi: 10.1002/PATH.4528.

[23] O. Kopper et al., “An organoid platform for ovarian cancer captures intra- and interpatient heterogeneity,” Nat Med, vol. 25, no. 5, pp. 838–849, May 2019, doi: 10.1038/s41591-019-0422-6.

[24] Y. Nanki et al., “Patient-derived ovarian cancer organoids capture the genomic profiles of primary tumours applicable for drug sensitivity and resistance testing,” Scientific Reports 2020 10:1, vol. 10, no. 1, pp. 1–11, Jul. 2020, doi: 10.1038/s41598-020-69488-9.

[25] M. Tao et al., “Developing patient-derived organoids to predict PARP inhibitor response and explore resistance overcoming strategies in ovarian cancer,” Pharmacol Res, vol. 179, p. 106232, May 2022, doi: 10.1016/J.PHRS.2022.106232.

[26] N. Maenhoudt et al., “Developing Organoids from Ovarian Cancer as Experimental and Preclinical Models,” Stem Cell Reports, vol. 14, no. 4, p. 717, Apr. 2020, doi: 10.1016/J.STEMCR.2020.03.004.

[27] N. V. Tzouras et al., “A Green Synthesis of Carbene-Metal-Amides (CMAs) and Carboline-Derived CMAs with Potent in vitro and ex vivo Anticancer Activity,” ChemMedChem, vol. 17, no. 13, p. e202200135, Jul. 2022, doi: 10.1002/CMDC.202200135.

[28] T. Scattolin et al., “The anticancer activity of an air-stable Pd(I)-NHC (NHC = N-heterocyclic carbene) dimer,” Chemical Communications, vol. 56, no. 81, pp. 12238–12241, Oct. 2020, doi: 10.1039/D0CC03883K.

[29] C. Granchi et al., “Design, synthesis and biological evaluation of second-generation benzoylpiperidine derivatives as reversible monoacylglycerol lipase (MAGL) inhibitors,” Eur J Med Chem, vol. 209, p. 112857, Jan. 2021, doi: 10.1016/J.EJMECH.2020.112857.

[30] T. Scattolin et al., “Palladium(II)-η3-Allyl Complexes Bearing N-Trifluoromethyl N-Heterocyclic Carbenes: A New Generation of Anticancer Agents that Restrain the Growth of High-Grade Serous Ovarian Cancer Tumoroids,” Chemistry – A European Journal, vol. 26, no. 51, pp. 11868–11876, Sep. 2020, doi: 10.1002/CHEM.202002199.

[31] G. E. Wensink et al., “Patient-derived organoids as a predictive biomarker for treatment response in cancer patients,” NPJ Precis Oncol, vol. 5, no. 1, Dec. 2021, doi: 10.1038/S41698-021-00168-1.

[32] M. Van De Wetering et al., “Prospective derivation of a living organoid biobank of colorectal cancer patients,” Cell, vol. 161, no. 4, pp. 933–945, May 2015, doi: 10.1016/j.cell.2015.03.053.

[33] W. Senkowski et al., “A platform for efficient establishment and drug-response profiling of high-grade serous ovarian cancer organoids,” Dev Cell, vol. 58, no. 12, pp. 1106-1121.e7, Jun. 2023, doi: 10.1016/j.devcel.2023.04.012.

[34] J. Jabs et al., “Screening drug effects in patient-derived cancer cells links organoid responses to genome alterations,” Mol Syst Biol, vol. 13, no. 11, Nov. 2017, doi: 10.15252/msb.20177697.

[35] C. Russo Spena et al., “Virtual screening identifies a PIN1 inhibitor with possible antiovarian cancer effects,” J Cell Physiol, vol. 234, no. 9, pp. 15708–15716, Sep. 2019, doi: 10.1002/jcp.28224.

[36] E. Cavarzerani, I. Caligiuri, M. Bartoletti, V. Canzonieri, and F. Rizzolio, “3D dynamic cultures of HGSOC organoids to model innovative and standard therapies,” Front Bioeng Biotechnol, vol. 11, 2023, doi: 10.3389/fbioe.2023.1135374.

[37] C. Russo Spena et al., “Liposomal delivery of a Pin1 inhibitor complexed with cyclodextrins as new therapy for high-grade serous ovarian cancer,” Journal of Controlled Release, vol. 281, pp. 1–10, Jul. 2018, doi: 10.1016/J.JCONREL.2018.04.055.

[38] C. E. Ford, B. Werner, N. F. Hacker, and K. Warton, “The untapped potential of ascites in ovarian cancer research and treatment,” British Journal of Cancer 2020 123:1, vol. 123, no. 1, pp. 9–16, May 2020, doi: 10.1038/s41416-020-0875-x.

[39] L. R. Hardy, A. Salvi, and J. E. Burdette, “UnPAXing the Divergent Roles of PAX2 and PAX8 in High-Grade Serous Ovarian Cancer,” Cancers 2018, Vol. 10, Page 262, vol. 10, no. 8, p. 262, Aug. 2018, doi: 10.3390/CANCERS10080262.

[40] S. A. Nik et al., “Upregulation of Wilms’ tumor gene 1 (WT1) in desmoid tumors,” Int J Cancer, vol. 114, no. 2, pp. 202–208, Mar. 2005, doi: 10.1002/IJC.20717.

[41] R. Koesters et al., “WT1 is a tumor-associated antigen in colon cancer that can be recognized by in vitro stimulated cytotoxic T cells,” Int J Cancer, vol. 109, no. 3, pp. 385–392, Apr. 2004, doi: 10.1002/IJC.11721.

[42] L. F. Sallum et al., “WT1, p53 and p16 expression in the diagnosis of low- and high-grade serous ovarian carcinomas and their relation to prognosis,” Oncotarget, vol. 9, no. 22, pp. 15818–15827, Mar. 2018, doi: 10.18632/ONCOTARGET.24530.

[43] W. Netinatsunthorn, J. Hanprasertpong, C. Dechsukhum, R. Leetanaporn, and A. Geater, “WT1 gene expression as a prognostic marker in advanced serous epithelial ovarian carcinoma: An immunohistochemical study,” BMC Cancer, vol. 6, no. 1, pp. 1–9, Apr. 2006, doi: 10.1186/1471-2407-6-90/FIGURES/3.

[44] D. Gupta and C. G. Lis, “Role of CA125 in predicting ovarian cancer survival - A review of the epidemiological literature,” J Ovarian Res, vol. 2, no. 1, pp. 1–20, Oct. 2009, doi: 10.1186/1757-2215-2-13/TABLES/9.

[45] M. E. L. van der Burg, F. B. Lammes, and J. Verweij, “CA125 in ovarian cancer,” 10.2217/17520363.1.4.513, vol. 40, no. 1–2, pp. 36–51, Dec. 2007, doi: 10.2217/17520363.1.4.513.

[46] M. Duffy et al., “CA125 in ovarian cancer: European Group on Tumor Markers guidelines for clinical use,” International Journal of Gynecologic Cancer, 2004, doi: 10.1136/IJGC-00009577-200509000-00001.

[47] P. Charkhchi, C. Cybulski, J. Gronwald, F. O. Wong, S. A. Narod, and M. R. Akbari, “CA125 and Ovarian Cancer: A Comprehensive Review,” Cancers 2020, Vol. 12, Page 3730, vol. 12, no. 12, p. 3730, Dec. 2020, doi: 10.3390/CANCERS12123730.

[48] N. Maenhoudt and H. Vankelecom, “Protocol for establishing organoids from human ovarian cancer biopsies,” STAR Protoc, vol. 2, no. 2, p. 100429, Jun. 2021, doi: 10.1016/J.XPRO.2021.100429.

[49] N. Maenhoudt et al., “Stem Cell Reports Article Developing Organoids from Ovarian Cancer as Experimental and Preclinical Models”, doi: 10.1016/j.stemcr.2020.03.004.

[50] G. Saorin et al., “Enhanced activity of a pluronic F127 formulated Pin1 inhibitor for ovarian cancer therapy,” J Drug Deliv Sci Technol, vol. 87, Sep. 2023, doi: 10.1016/j.jddst.2023.104718.

[51] E. A. Brooks, S. Galarza, M. F. Gencoglu, R. Chase Cornelison, J. M. Munson, and S. R. Peyton, “Applicability of drug response metrics for cancer studies using biomaterials,” Philosophical Transactions of the Royal Society B: Biological Sciences, vol. 374, no. 1779. Royal Society Publishing, Aug. 19, 2019. doi: 10.1098/rstb.2018.0226.

