## Supplemental Figure S1 for "Longitudinal Prediction of Drug Response in High-Grade Serous Ovarian Cancer Organoid Cultures Aligning with Clinical Responses"

### SUPPLEMENTARY FIGURES

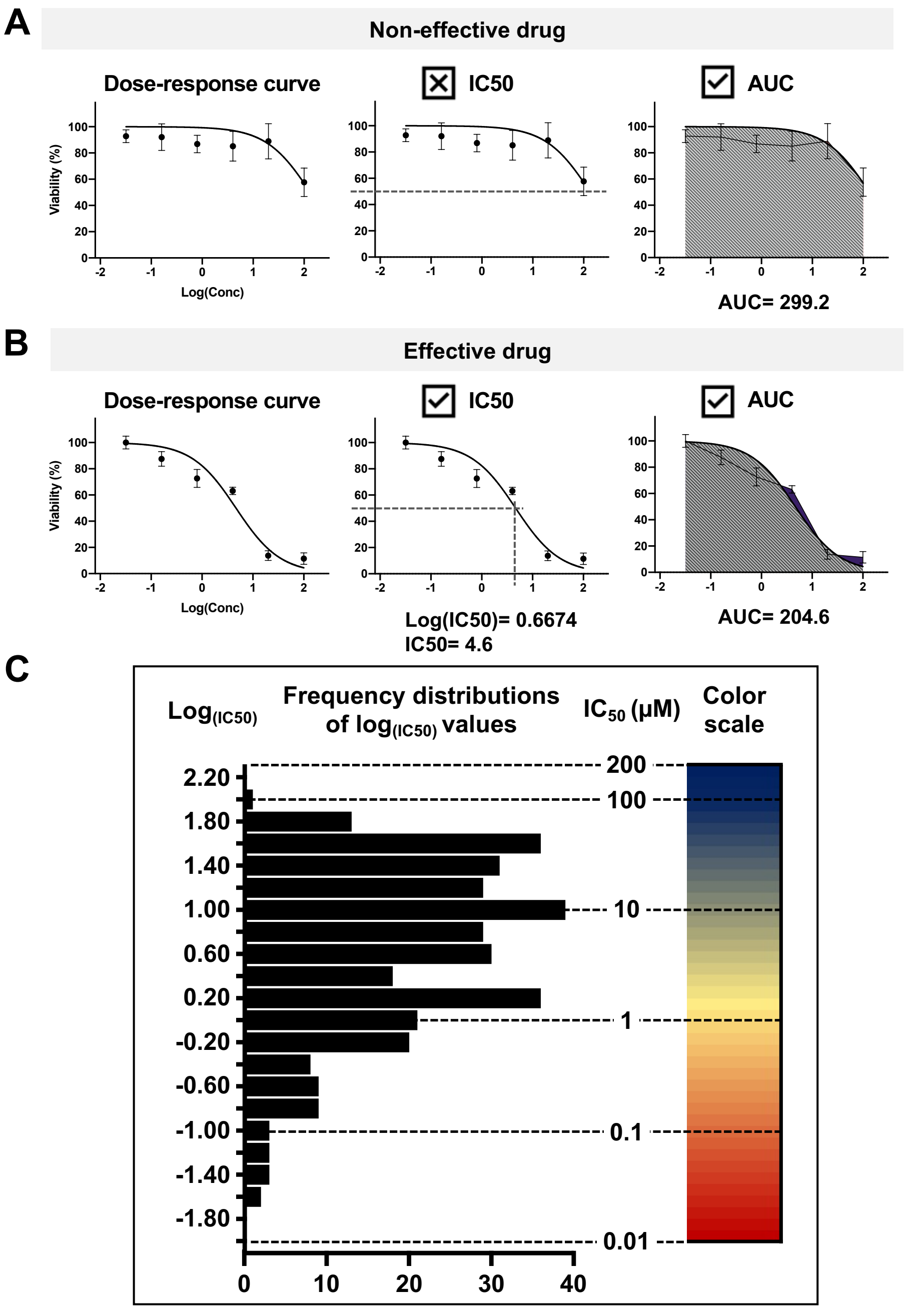


***Figure S1. Drug response interpretation:*** *The IC50 represents the drug concentration where the response is reduced by half and the AUC represents the area under the dose–response curve. The y-axis shows the normalized growth rate. Drugs are considered effective (B) when an IC50 value can be calculated, that is, when the drug decreases viability by more than 50% at the maximum dose. Conversely, drugs that do not reduce viability by at least 50% at the maximum dosage are considered non-effective (A). Regardless of whether a drug is effective or not, the AUC can be calculated. (C) For all effective drugs for which an IC50 value can be determined, a color gradient ranging from red to blue is assigned, where red indicates the most effective drugs, and blue/green indicates the least effective, meaning those with the higher IC50 values.*
